# Dynamic proteomics of HSV1 infection reveals molecular events that govern non-stochastic infection outcomes

**DOI:** 10.1101/081653

**Authors:** Nir Drayman, Omer Karin, Avi Mayo, Tamar Danon, Lev Shapira, Oren Kobiler, Uri Alon

## Abstract

Viral infection is usually studied at the level of cell populations, averaging over hundreds of thousands of individual cells. Moreover, measurements are typically done by analyzing a few time points along the infection process. While informative, such measurements are limited in addressing how cell variability affects infection outcome. Here we employ dynamic proteomics to study virus-host interactions, using the human pathogen Herpes Simplex virus 1 as a model. We tracked >50,000 individual cells as they respond to HSV1 infection, allowing us to model infection kinetics and link infection outcome (productive or not) with the cell state at the time of initial infection. We find that single cells differ in their preexisting susceptibility to HSV1, and that this is partially mediated by their cell-cycle position. We also identify specific changes in protein levels and localization in infected cells, attesting to the power of the dynamic proteomics approach for studying virus-host interactions.

## INTRODUCTION

Viral infection is a heterogeneous process. One example is the variation in the number of viral progeny produced by individual cells, which spans several orders of magnitude, as first described for bacteriophages in the 1940's (Delbrück, 1945). Several recent studies found similar variability in mammalian viruses (Zhu et al., 2009; Timm and Yin, 2012; Schulte and Andino, 2014; Combe et al., 2015; Heldt et al., 2015).

Viral infection course can also vary, between a lytic and a lysogenic (or latent) cycle. For HIV1, this decision is stochastic and governed by the initial fluctuations in the number of viral Tat proteins, leading to a bi-stable decision (Weinberger et al., 2005; Singh and Weinberger, 2009; Razooky et al., 2015). In contrast, the decision between lysogeny and lytic infection of phage Lambda seems to be more dependent on the host cell, with smaller bacteria more likely to undergo lytic infection (St-Pierre and Endy, 2008).

Variability also exists in infection outcome, when some cells in a population become successfully infected, while others resist the infection. One known source of this variability is cell-extrinsic, emanating from the random distribution of the number of viruses that individual cells encounter. This distribution explains why, on the population level, the percentage of infected cells is governed by Poisson's law according to the number of infectious units per cell - the multiplicity of infection (moi) (Parker, 1938; Smith, 1968).

In addition, infection outcome might also be influenced from the host cellular state at the time of virus adsorption. Such cell-intrinsic differences include the cell-cycle stage and cell-to-cell variability in protein levels and activities that have been studied in other contexts (Elowitz et al., 2002; Cohen et al., 2008; Tay et al., 2010; Loewer and Lahav, 2011; Kellogg and Tay, 2015).

Supporting a possible role for the host cellular state in determining infection outcome, Pelkmans et al. performed studies in which cells were infected with different viruses, fixed several hours later and immuno-stained for viral proteins. Using a machine learning approach, they found that infected cells differ from non-infected cells (Snijder et al., 2009, 2012). However, as cells were not imaged at the time of virus adsorption, it is unclear whether the observed differences between the cells are the cause or the consequence of viral infection success.

There is a lack of experiments that directly address the question of determinism in viral infection, which requires a system that follows individual cells from the time of virus adsorption to the onset of viral gene expression (distinguishing successful from failed infections).

To address this question, we use the dynamic proteomics approach (Sigal et al., 2006a, 2006b, 2007; Geva-Zatorsky et al., 2010; Eden et al., 2011; Farkash-Amar et al., 2014) that relies on a library of human cell-line clones, each expressing a unique YFP-tagged host protein from its native chromosomal location. All the clones also express mCherry-tagged markers that are used for automated segmentation and tracking of the cells. We follow tens of thousands of individual cells from ~400 clones throughout the process of infection by the human pathogen Herpes Simplex virus 1 (HSV1). The virus expresses mTurquoise2, a bright variant of CFP (Goedhart et al., 2012), allowing the monitoring of successful infection in real-time. We start imaging at the time of viral adsorption, monitoring infection kinetics, cellular proteins level and localization as well as the cellular environment through time-lapse fluorescence microscopy (Fig. 1A).

**Figure 1.**
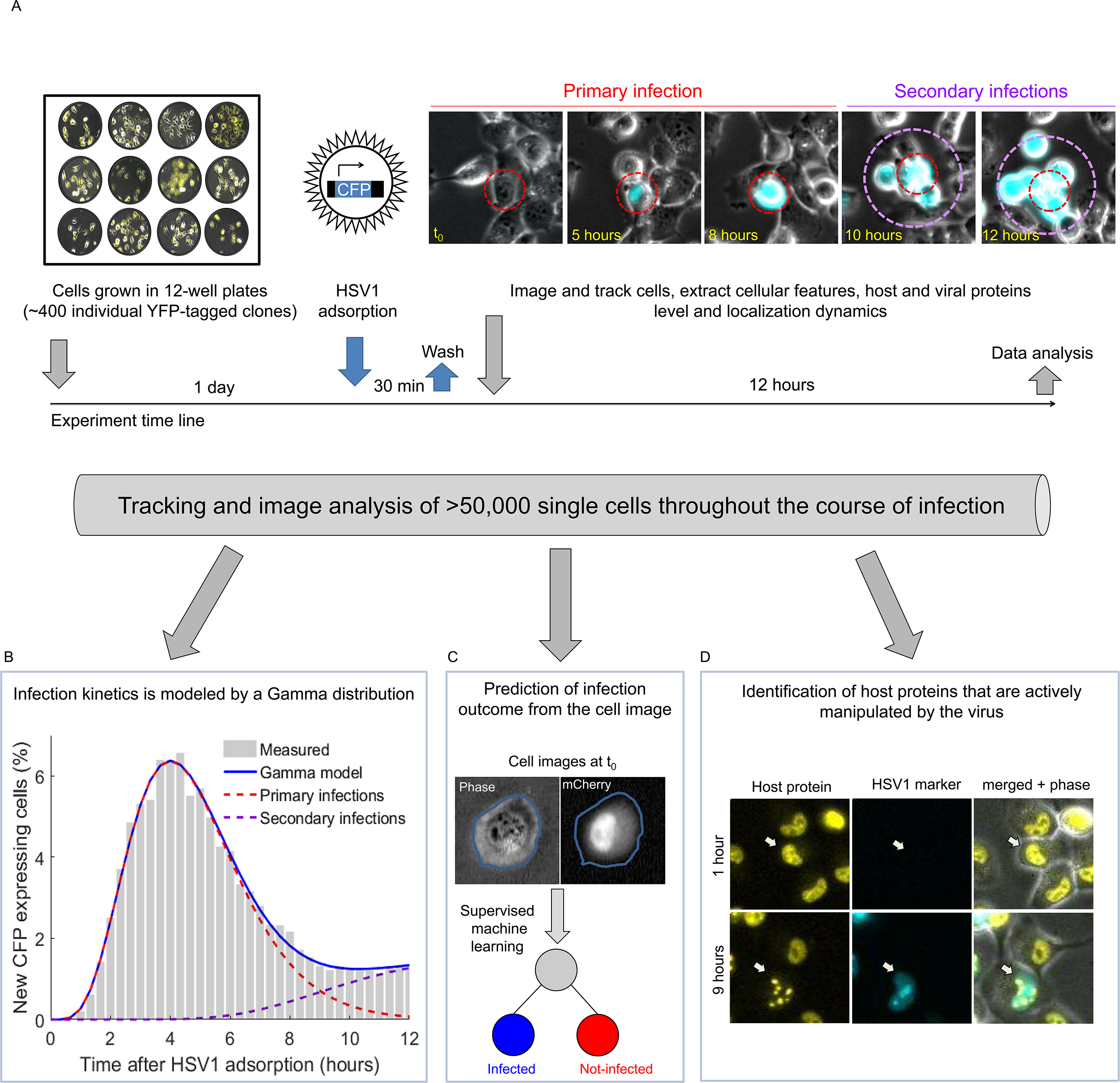
Dynamic proteomics to study virus-host interactions in single cells over time. **A.** Schematic representation of the screen. A CFP-expressing HSV1 was allowed to adsorb to clones seeded in 12-well plates for 30 minutes, washed out and cells subsequently imaged every 20 minutes for 12 hour. Overall, more than 50,000 single cells were followed, from ~400 different YFP-expressing clones **B.** Model for de-mixing primary and secondary infections. Shown are the measured lag-times between virus adsorption and CFP expression (gray bars), the fitted model (blue line) which is based on primary (red dashed line) and secondary (purple dashed line) infections. See Supplementary Fig. 2 for more information. **C.** A supervised machine learning approach was used to predict infection outcome from the cell images at the time of virus adsorption. See Fig. 2 for more information. **D.** Specific changes in host proteins levels and localization upon HSV1 infection were studied. Shown is an example of a localization change in cells infected by HSV1. See figures 5 and 6 for more information.

This dataset allows us to probe multiple aspects of the virus-host interaction at the single cell level (Fig. 1B-D). By measuring and modeling the distribution of lag-times between virus adsorption and viral gene expression in the population (Fig. 1B), we discover that infection kinetics are well described by a Gamma distribution. We next ask if there is a role for the host cellular state in determining infection outcome. We find that 20 image-analysis features extracted from cells at the time of adsorption can be used to predict the outcome of viral infections nine hours later (Fig. 1C). These features include the cell's velocity, cell-cycle stage and nuclear area, among others. We further find that this cell-intrinsic difference in susceptibility to HSV1 infection is pre-existing in the population, as infection outcome can be determined even when using cell images taken 24 hours prior to the cells encountering the virus. Finally, we analyze the effect of HSV1 infection on the host proteins (Fig. 1D). Of the ~400 host proteins studied, two (Geminin and RFX7) showed a significant difference in their concentration at the time of virus adsorption between cells that will become successfully infected and those that will not. Two others (SUMO2 and RPAP3) were degraded specifically in infected cells, and another two (SLTM and YTHDC1) changed their localization upon infection.

## RESULTS

### Dynamic proteomics of human cells following HSV1 infection

We infected ~400 clones from our library with a CFP-expressing HSV1 at a multiplicity of infection (moi) of 0.5. CFP expression correlates with expression of the viral immediate-early protein ICP4 (Supplementary Fig. 1), which is required for the progression of successful infection. This design allowed us to compare successfully-infected and non-infected cells side by side. Note that in this cell line, the moi used is equivalent to ~50 virus particles per cell, such that all cells in the culture have likely encountered viruses. H1299 cells are fully permissive for HSV1 infection, as evident by the spread of infection from primary infected cells to produce secondary infections (Fig. 1A, Supplementary Movie 1).

Using custom software we tracked tens of thousands of individual cells for 12 hours after HSV1 adsorption at a time resolution of 20 minutes, extracting features such as the cell's position, shape and size, as well as the level and sub-cellular localization of the different fluorescent proteins. Since we continuously monitored the cells we could identify the point at which each infected cell began to express CFP. We refer to this time delay between viral adsorption and initial expression of viral encoded proteins as infection lag time. We found that the lag time is variable among individual cells with a mean time of 5.9±2.6 hours and a CV of 44% (Fig. 1B).

Since HSV1 undergoes productive infection in the cells, the distribution of lag times captures both primary infections and subsequent secondary infections (Fig. 1A,B). To determine the kinetics of infections we modeled infection kinetics as the sum of primary and secondary infections (Fig. 1B and Supplementary Fig. 2). We fitted our model with two-parameter distributions (Normal, Log-normal, Weibull and Gamma), estimating for each distribution the best fitted parameters. We found that the lag-time distribution is best described by a Gamma distribution with a shape factor of 6 and a rate parameter of 1.25 (Supplementary Fig. 2 and Methods).

In addition to the theoretical interest in modeling infection kinetics, this model also allowed us to determine a cut-off point between primary and secondary infections, which is 9 hours post virus adsorption. Here we are only interested in the analysis of the primary infections, which more closely resembles the initial infection of a human host. In all subsequent analyses, we considered cells as successfully infected if their initial CFP expression time was below 9 hours and as non-infected if they remained CFP negative for the entire 12 hours. Cells whose CFP expression time was between 9-12 hours were removed from the analyses.

As we started our time-lapse recording from viral adsorption, we could not directly observe the cell's position along the cell-cycle (based on its previous mitosis). To circumvent this, we determined the cell-cycle stage from still images, similarly to what was previously done by others (Kafri et al., 2013; Gut et al., 2015; Blasi et al., 2016) (See Methods and Supplementary Fig. 3 for more information).

### HSV1 infection outcome is largely determined by the host cellular state at the time of adsorption

Overall, we tracked >52,000 single cells from ~400 different clones for 12 hours after HSV1 adsorption. We determined primary infection outcome (successful or not) based on the cell's CFP levels at nine hours post adsorption. Of these cells, 22,182 were successfully infected (CFP positive) and 29,993 were not (CFP negative). The percentage of infected cells varied among different clones and was 35±13% CFP positive cells.

To test whether infection outcome is dependent on the cellular state at the time of virus adsorption (time zero) we employed a supervised machine learning approach. If infection outcome depends on the cellular state, then features extracted from the cell image should contain information regarding the future outcome of the virus-host encounter. We therefore trained a decision-tree based classifier to predict infection outcome from features extracted from the cell image at the first two frames of the movie (Fig. 2A and Methods). We extracted standard image-analysis features (listed in Supplementary Table 1) of cell morphology (53 features such as cell size and roundness), texture (320 features such as homogeneity, contrast and dissimilarity), velocity, local cell density and cell-cycle stage.

**Figure 2.**
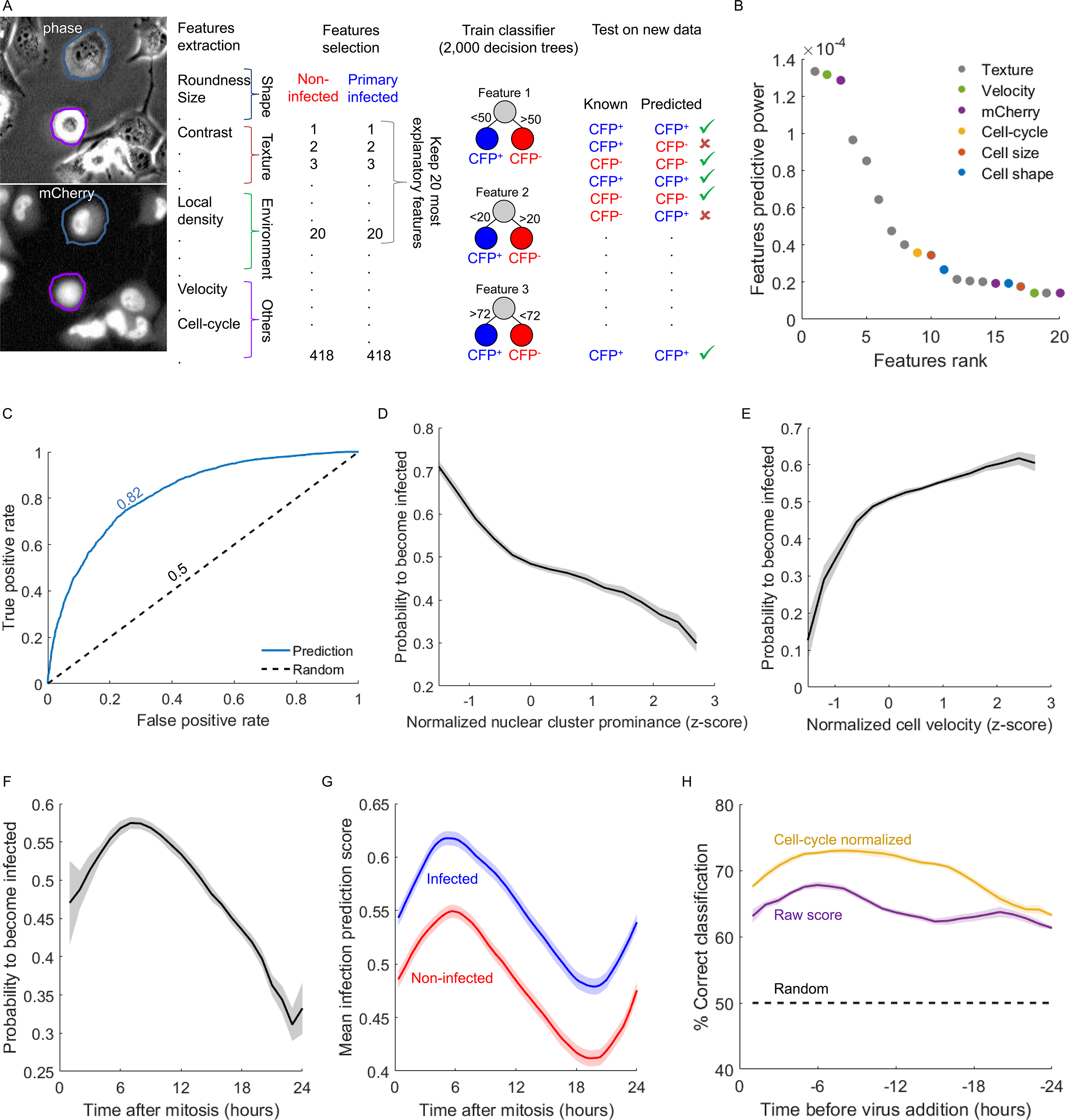
HSV1 infection outcome can be predicted from images of the cells at the time of adsorption. **A.** Schematic representation of the machine learning approach; First, image-analysis features were extracted for each cell from the phase and mCherry channels. Second, of the 418 features calculated for each cell, we chose the top 20 most explanatory features to continue with. The full list of features with their relative explanatory power is listed in Supplementary Table 1. We trained a supervised machine learning classifier to best discriminate cells that will becomes successfully infected from those that will not and tested its performance on a separate test set. **B.** The top 20 ranking image-analysis features that were used for predicting infection outcome. For convenience we color coded the features according to the next groups: Texture (gray) - all textural features, extracted from either the phase or mCherry channel. Velocity (green) normalized (feature ranked #2) or raw (#18). mCherry (Purpule) - normalized mCherry concentration (#3), raw nuclear mCherry level (#15), normalized mCherry level (#20). Cell-cycle position at the time of infection (yellow). Cell size (red) - nuclear area (#10), change in normalized cell width (#17). Cell Shape (blue) - normalized cell roundness (#11), cell eccentricity (#16). **C.** ROC curves for the trained classifier (blue line) and a random prediction (dashed black line) The area under the curves are noted in corresponding colors. **D,E,F**. Probability for cells to become infected, based on three different cellular features at time zero. D cluster prominence (textural feature), E - cell velocity and F - cell-cycle position. **G.** Them mean classifier score in cells that will become infected (blue) or nor-infected (red) over 24 hours, aligned by their cell-cycle position. The figure shows the clear cell-cycle dependency for infection outcome, as well as the cell-cycle independent component, as the two curves never cross each other. **H.** Correct classification can also be done using images of cells prior to the addition of the virus. A second, smaller dataset was constructed in which cells were first imaged for 24 hours and then infected. Shown are the classifier performances based on the raw predictor score (purple line) or after normalizing by the cell-cycle stage (yellow line).

We trained the classifier on 23,780 cells, using the 20 most explanatory features (Fig. 2B, Supplementary Table 1). The features used include textural features as well as the cell's velocity, mCherry concentration, cell-cycle stage, nuclear area and cell morphology. The classifier outputs the probability of a cell to become infected, ranging from 0-1, and we refer to this output hereafter as the classifier score. We classified cells based on the classifier score, using a threshold of 0.5 to assign cells to the infected or non-infected groups. We tested the classifier performance on a dataset of 8,108 cells from clones not used in the training step and found that it correctly predicted infection outcome in 75% of these cells with an area under the curve (AUC) of 0.82 (Fig. 2C).

We next analyzed the contribution of the different features in predicting infection outcome. The most predictive feature was the nuclear cluster prominence, a measurement of textural asymmetry (Unser, 1986). Cells with low cluster prominence were more likely to become successfully infected, and the probability for infection decreased as the cluster prominence increased (Fig. 2D).

The second most predictive feature was the cell's velocity at the time of virus adsorption (Fig. 2E). While infection probability continues to increase as the cell movement increases, the most pronounced effect was seen in cells with low velocity (z-score<0) which were more resistant to viral infection. Infection probability dependence on the top 10 features can be found in Supplementary Fig. 4.

The cell-cycle stage was ranked ninth among these features (Fig. 2B). Infection probability showed a non-monotonic relation to the cell-cycle stage, peaking around six hours after mitosis and then decreasing as the cells progress through the cell cycle (Fig. 2F). We tested whether the cell-cycle effect is independent from that of other features by comparing the mean classifier score of cells that will become infected and those that will not over 24 hours, after aligning them to the same stage of the cell cycle (Fig. 2E). We observed that the classifier score is higher in cells that will become infected throughout the cell-cycle, implying that the cell-cycle effect is at least partially independent from that of other features, such as the cell velocity and texture that were described above.

The success of a machine learning algorithm in predicting the success of viral infection in individual cells suggests that the outcome of infection is not intrinsically stochastic but rather depends on the cellular state at the time of infection.

### Cellular susceptibility is pre-existing in the population prior to encountering the virus

Since our time-lapse recordings started approximately 45 minutes after HSV1 adsorption to the cells, we wanted to verify that the classifier was not influenced by a rapid response of the cells to the infection. To address this, we performed longer time-lapse movies in which we first imaged the cells unperturbed for 24 hours before adding the virus. We tracked 124 infected cells and 99 non-infected cells.

We analyzed the classifier's performance when using cell images up to 24 hours prior to the addition of the virus. We did so by using either the raw score given by the classifier (Fig. 2H ,purple line), or after normalizing according to the cell-cycle effect, as the cells move along the cell-cycle phases during these 24 hours (Fig. 2H ,orange line). Both analyses showed that susceptibility to HSV1 infection is long-lasting, with 61-63% correct classification achieved even when using images from 24 hours prior to the cells encountering the virus. This is especially apparent when controlling for the cell-cycle effect, with cell images taken 17 hours prior to virus adsorption resulting in ~70% correct classification (Fig. 2H).

To quantify the life time of infection susceptibility we calculated the mixing time of the predictor output (Sigal et al., 2006b). This is done by computing the auto-correlation of the predictor output over time. We found that the mixing time (the time it takes for the auto-correlation to decay to 0.5) is around 6 hours when using the raw data and 10 hours when controlling for the cell-cycle effect (Supplementary Fig. 5).

Taken together, our findings suggest that the cellular state that underlies susceptibility to HSV1 infection is composed of: (1) a faster changing, cell-cycle dependent component and (2) a more stable, cell-cycle independent component.

### Cells in the early part of the cell-cycle are more susceptible to HSV1 infection

To better understand the molecular mechanism that underlies the variability in infection outcome among individual cells, we looked for proteins whose concentration significantly differ between cells that will become infected and those that will not at time zero. Of the ~400 proteins screened, we identified two such proteins - Geminin and RFX7 (Fig. 3A). On average, Geminin concertation was 40% lower in cells that will become infected and RFX7 concentration was 37% lower (Fig. 3B). The difference in their concentration was mainly observed at time zero, and disappeared later in the infection (Fig. 3C,D).

**Figure 3.**
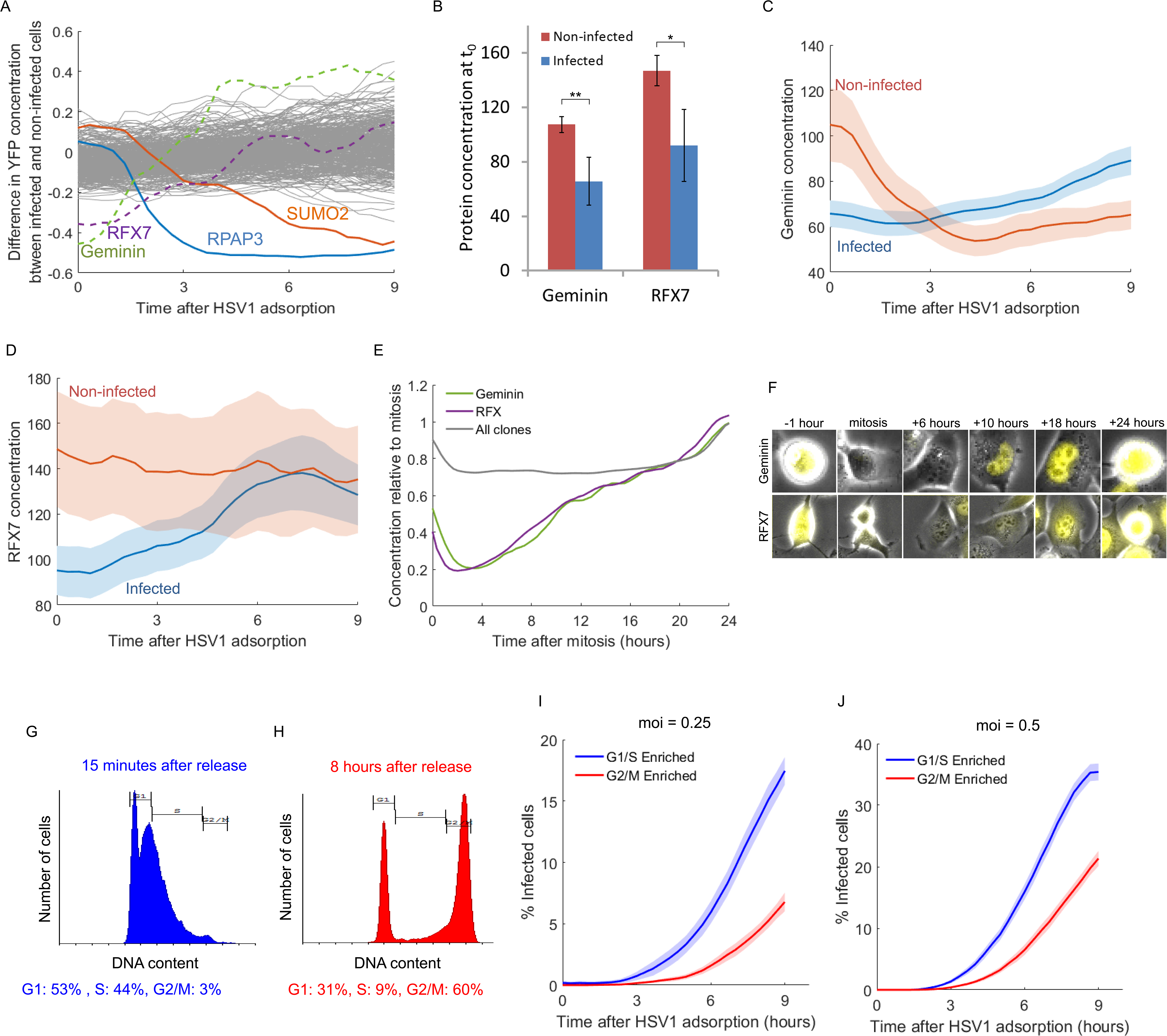
Cells in the early part of the cell-cycle are more susceptible to HSV1 infection. **A.** The fraction difference in YFP concentration between infected and non-infected cells over the first nine hour after adsorption. The majority of the clones (grey lines) showed differences in the range of ±0.2 (20%), which are within experimental error. Four clones are highlighted. Geminin (dashed green line) and RFX7 (dashed purple line) concentrations are lower in cells that will become infected at time zero. SUMO2 (red line) and RPAP3 (blue line) concertation decrease over time, specifically in cells that will become infected. **B.** Geminin and RFX7 concentration (mean±s.e.m) in cells that will become infected (blue bars) or not-infected (red bars) at time zero. * p-value<0.05, ** p-value<0.01. Both were calculated based on a one-tailed t-test. **C,D.** Geminin (C) and RFX7 (D) concentration (mean±s.e.m) over time in cells that will become infected (blue lines) or not-infected (red lines). **E.** Geminin (green line) and RFX7 (purple line) concentration are cell-cycle dependent. The gray line depicts the median behavior of all other clones in the screen. **F.** Representing images of cells expressing YFP-Geminin (top row) or YFP-RFX7 (bottom row) at different times post-mitosis. **G,H.** Distribution of DNA content in cells 15 minutes (G) and 8 hours (H) after release from double-thymidine block. **I,J.** Accumulation of infected cells (CFP^+^) as a function of time for cells enriched for the G1 and S (blue lines) or G2/M (red lines) stages, infected with HSV1 at an moi of 0.25 (I) or 0.5 (J). mean±s.e.m from six positions for each moi.

Geminin levels are known to be correlated with the cell-cycle stage, as Geminin is a substrate of the Anaphase Promoting Complex (McGarry and Kirschner, 1998). RFX7 is a member of a transcription factor family that binds the X-box motif, which are important for the regulation of immune genes such as MHC class II (Fontes et al., 1997). We found that both Geminin and RFX7 show a similar cell-cycle related concentration profile (Fig 3E). Both are rapidly degraded following mitosis, with their concentration rising slowly towards the next mitosis (Fig. 3F).

The lower concentrations of Geminin and RFX7 in cells that will become infected is an independent indication that cells in the earlier part of the cell-cycle are more susceptible to HSV1 infection, in agreement with the results obtained from the machine learning approach described above.

To further experimentally test the effect of the cell cycle on infection outcome we used the double thymidine block protocol, which synchronizes cells to the G1/S checkpoint (Bootsma et al., 1964). We infected cells either 15 minutes after releasing from the block (Fig. 3G) or 8 hours after release, where the majority of cells are in the G2/M stages (Fig. 3H). We found that cells infected during the G1 and early S phases were 2-3 fold more susceptible to HSV1 infection than cells infected as the G2 and M phases, in two multiplicities of infection (Fig. 3I,J).

Taken together, our results clearly demonstrate a deterministic role for the cell-cycle stage in determining HSV1 infection outcome among individual cells.

### Infection kinetics are affected by position along the cell cycle

After considering the effect of the cell cycle on infection outcome, we next considered its effect on infection kinetics. We divided the cells in our dataset into early (G1 and early S) and late (late S, G2 and M) stages and measured their distribution of infection lag times (Fig. 4A). We observed a clear effect of the cell-cycle on infection kinetics with cells at the early cell cycle stages infected 20% faster than cells at the late stages (Fig. 4A,B).

**Figure 4.**
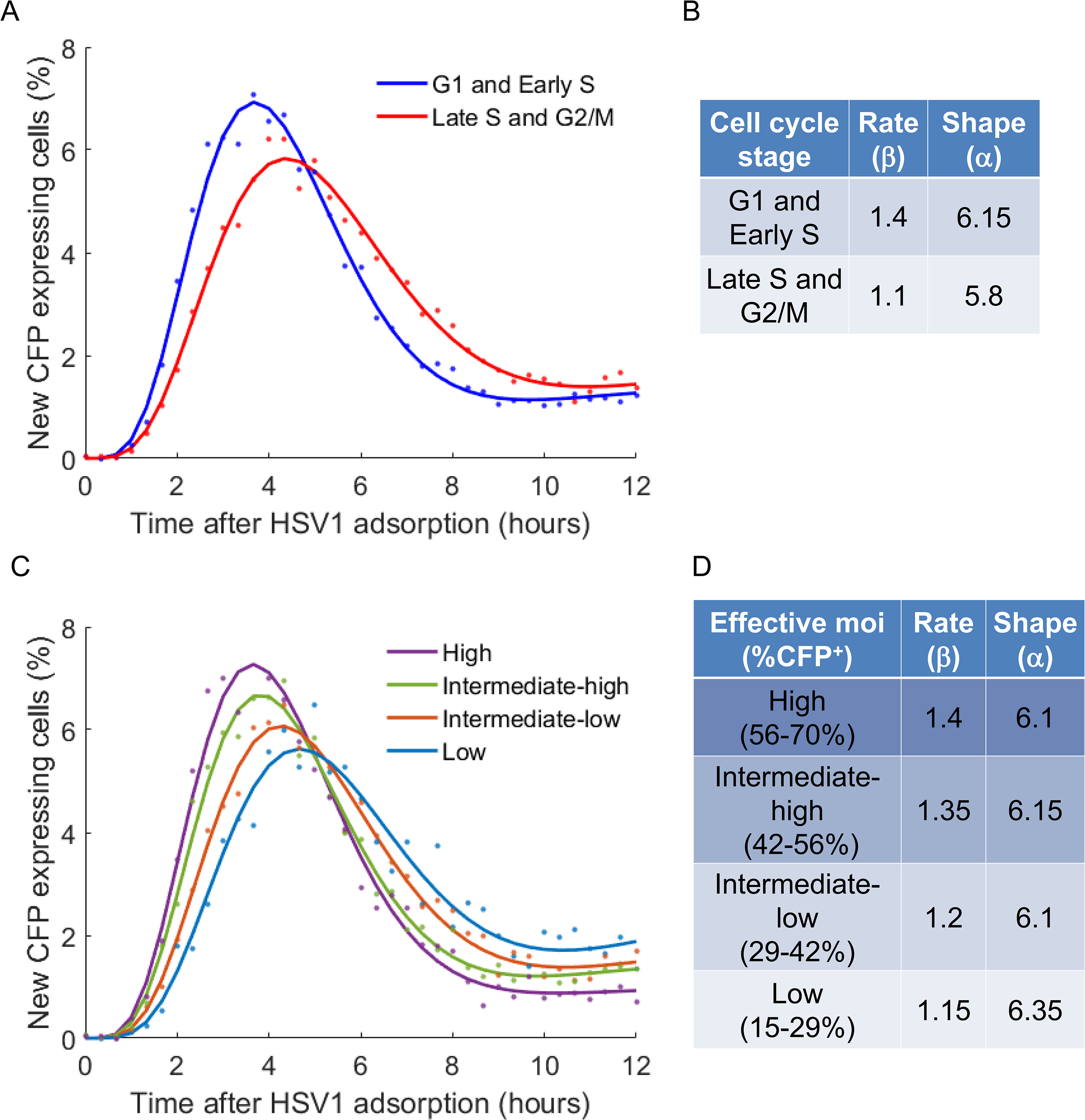
The cell-cycle stage of the host cell impact HSV1 kinetics. **A.** Data points (circles) and models (lines) fitted to cells infected at the early part of the cell-cycle (G1 and early S) or the late part of the cell-cycle (late S and G2/M). Cells in the G1 and early S stages show a faster kinetics with shorter lag times. **B.** Best estimated parameters for the Gamma distributions of primary infection kinetics shown in A. Cells at the later stages of the cell-cycle (Late S and G2/M) showed a 21% decrease in the rate parameter β. The shape parameter α was largely unaffected, showing a 6% difference. **C.** Data points (circles) and models (lines) fitted to cells binned by their effective moi. Infection kinetics is faster as the effective moi increases. **D.** Best estimated parameters for the Gamma distributions of primary infection kinetics shown in C. The rate parameter β increases as the effective moi increases, while the shape parameter α remains largely unaffected, with a slight increase (4%) at the lowest moi.

We compared the effect of the cell-cycle stage (a cell-intrinsic variability source) to that of the effective moi (a cell-extrinsic variability source). As explained above, although all experiments were conducted in the same moi of 0.5, a large variation in the percentage of infected cells was observed. We used this experimental variation to bin different time-lapse movies into four equally-spaced categories of effective moi, from low (15-29% CFP positive cells) to high (5670% CPF positive). The distributions of lag times and the fitted models for different effective moi are shown in Fig. 4C,D.

We found that cells infected during G1 and early S stages show similar kinetics to cells in the highest effective moi. Cells infected during late S and G2/M stages showed kinetics similar to those in the lowest effective moi. Thus, the cell-cycle stage at the time of virus adsorption has an effect of similar magnitude to that of changing the effective moi by a factor of ~3. Furthermore, both the cell-cycle stage and the moi scale the kinetics of the infection by changing the rate parameter of the Gamma distribution (β) without affecting the shape parameter (α).

We conclude that HSV1 infection kinetics is affected by the cell-cycle stage of the host cell at the time of viral adsorption.

### HSV1 infection causes a sharp decline in SUMO2 and RPAP3 concentrations

Having considered the effect of the cellular state on infection outcome, we next turned to study the effect of viral infection on the host cell. To our surprise, the majority of the ~400 host proteins studied did not show significant differences in concentration between infected and non-infected cells (Fig. 3A).

However two proteins, SUMO2 and RPAP3, showed reduced protein concentrations following adsorption in cells that eventually became infected (Fig. 3A). SUMO2 is a ubiquitin homolog that can be covalently attached to cellular proteins. Indeed, a decrease in SUMO2 levels upon HSV1 infection has been previously reported (Sahin et al., 2014; Sloan et al., 2015). SUMO2 concentration began dropping approximately two hours after HSV1 infection and continued so for the next nine hours (Fig. 5A,C and Supplementary Movie 2).

**Figure 5.**
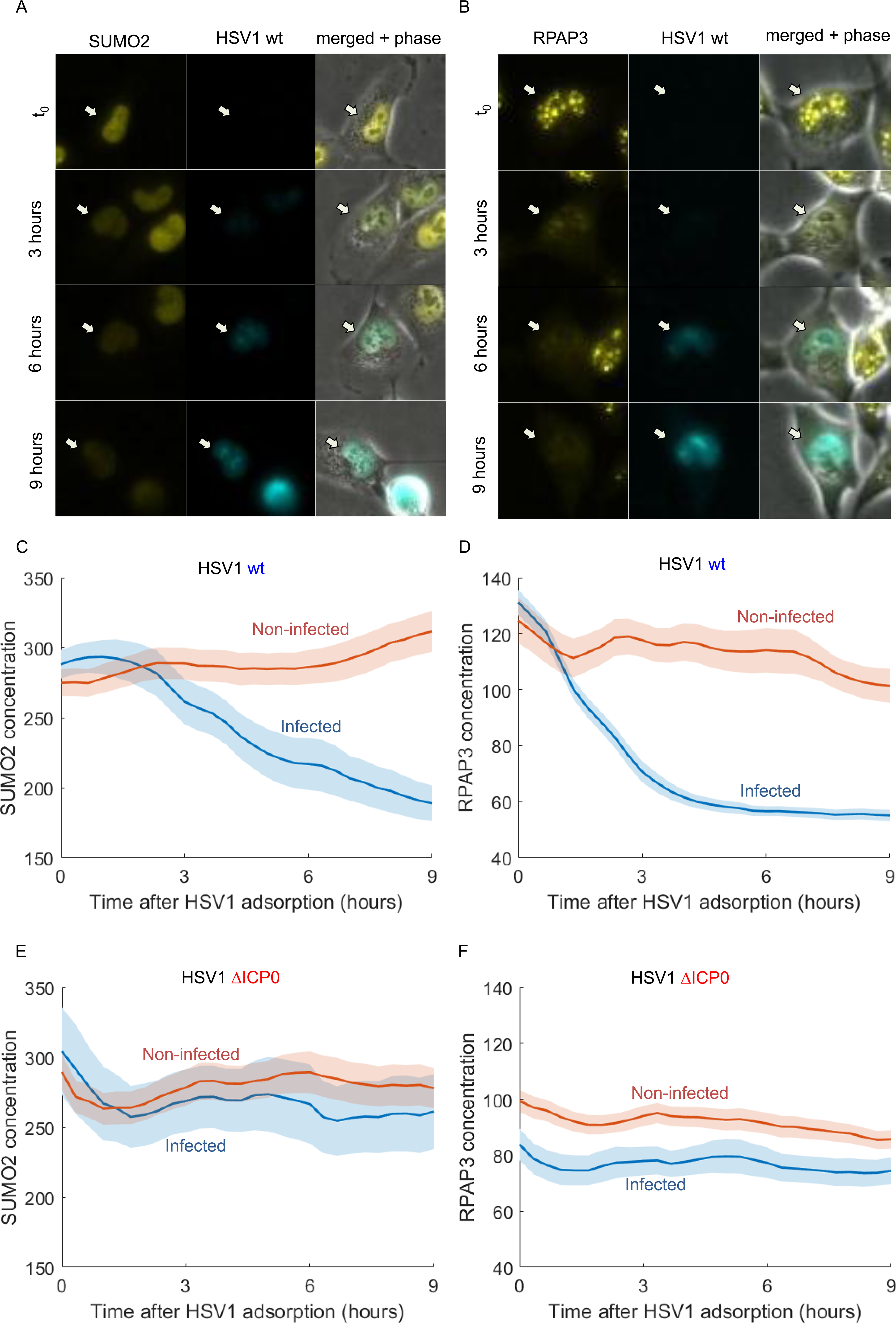
SUMO2 and RPAP3 degradation upon HSV1 infection is facilitated by ICP0. A,B. Images of representing cells from time-lapse movies of SUMO2 (A) and RPAP3 (B). Shown are the YFP channel, CFP channel and a merged image including the phase channel at 0-9 hours post wild-type HSV1adsorption. **C,D.** mean±s.e.m of SUMO2 (C) and RPAP3 (D) concentrations in non-infected (red) and infected (blue) cells following wild-type HSV1 adsorption. **E,F.** mean±s.e.m of SUMO2 (E) and RPAP3 (F) concentrations in non-infected (red) and infected (blue) cells following infection by a mutant HSV1 that does not express ICP0

RPAP3 (RNA polymerase II-associated protein 3) has not been previously characterized in the context of viral infection. In non-infected cells it showed distinct foci in the nucleus, which may represent transcriptional complexes on the cellular DNA (Fig. 5B). In HSV1 infected cells RPAP3 levels rapidly dropped, beginning at time zero and preceding CFP expression (Fig. 5B,D and Supplementary Movie 3). The degradation of RPAP3 might relate to the previously observed changes in RNA-polymerase II dependent transcription following HSV1 infection (Wagner and Roizman, 1969; Rice et al., 1994; Spencer et al., 1997; Jenkins and Spencer, 2001; Abrisch et al., 2015).

As the degradation of SUMO2 and RPAP3 seem to be very rapid, it is probably mediated by one of HSV1 immediate-early proteins. The most likely candidate for this is the virus encoded E3-ubiquitin ligase - ICP0. We tested this by infecting the cells with a mutant virus that does not express ICP0 and expresses a CFP reporter. Indeed, cells successfully infected by the ICP0 null mutant did not show degradation of either SUMO2 or RPAP3 (Fig. 5E,F and Supplementary Movies 4,5).

### HSV1 infection causes re-distribution of SLTM and YTHDC1

One unique feature of live cell microscopy is the ability to observe changes in the localization of tagged proteins. We studied these changes by looking at the nuclear/cytoplasm ratio of the proteins, and on their coefficient of variance (CV) in the nucleus and cytoplasm (which indicate how dispersed is the protein in these compartments). We did not observe any nucleus/cytoplasm trafficking nor changes in cytoplasmic proteins CV. We found that the CV of two nuclear proteins (SLTM and YTHDC1) increased specifically in successfully infected cells (Fig. 6A,B,C). Both SLTM and YTHDC1 participate in splicing of mRNA (Nayler et al., 1998; Xiao et al., 2016). The increase in the nuclear CV is a result of re-distribution of these proteins upon infection, from being equally diffused around the nucleus to forming large foci (Fig. 6 D,E and Supplementary Movies 6,7).

**Figure 6.**
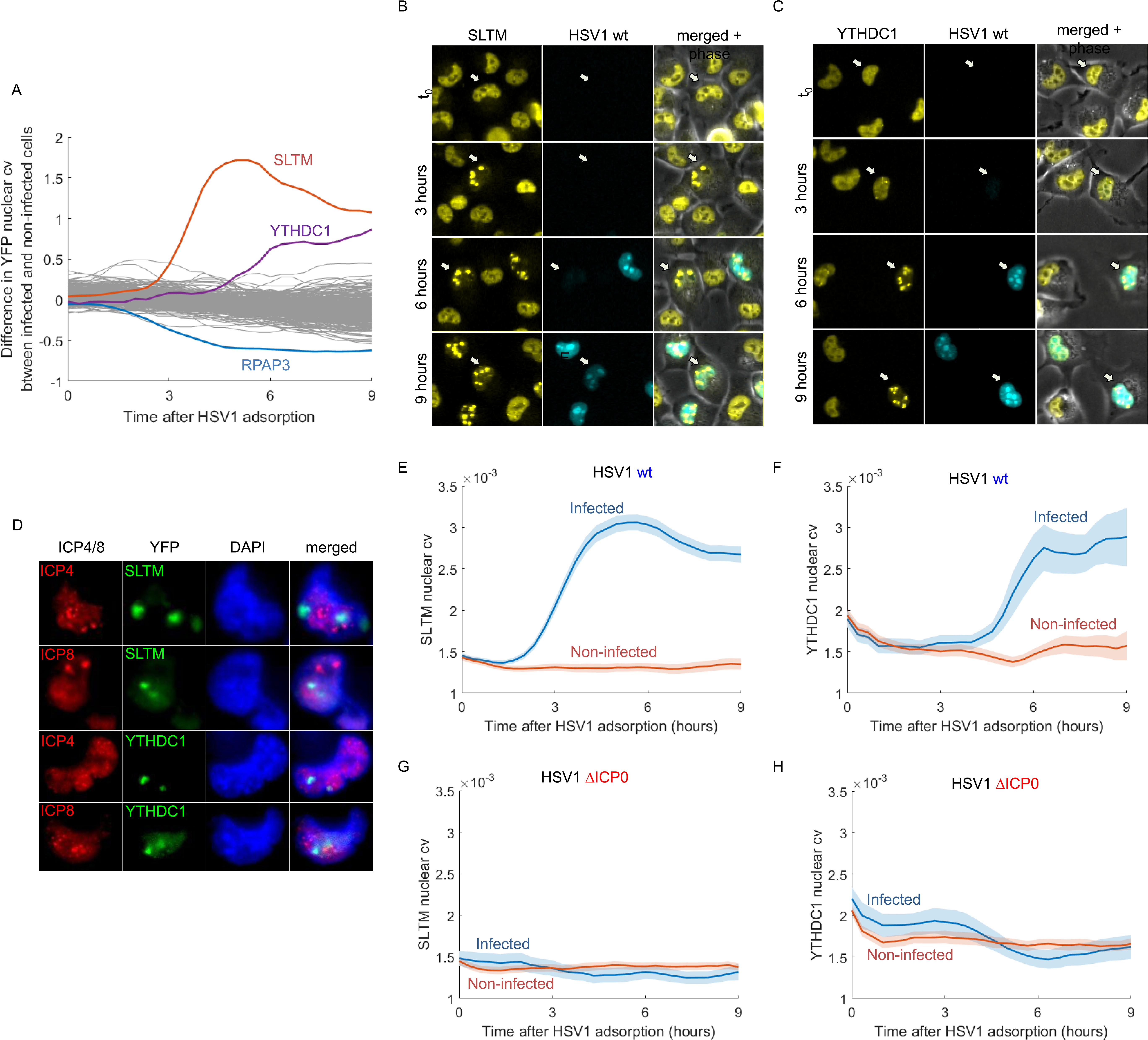
SLTM and YTHDC1 localization change upon HSV1 infection is facilitated by ICP0. **A.** The fraction difference in YFP nuclear cv between infected and non-infected cells over the first nine hour after adsorption. The majority of the clones (grey lines) did not show significant changes. Three clones are highlighted; SLTM (red line) and YTHDCH1 (purple line) which show an increase in nuclear cv and RPAP3 (blue line) which shows a decrease in nuclear cv. **B,C.** Images of representing cells from time-lapse movies of SLTM (B) and YTHDC1 (D). Shown are the YFP channel, CFP channel and a merged image including the phase channel at 0-9 hours post wild-type HSV1adsorption. **D.** Cells infected by wild-type HSV1 were fixed and stained for ICP4 or ICP8 at six hours post adsorption and imaged using a X100 magnification lens. Shown are representative images of nuclear foci formed by SLTM (top two rows) or YTHDC1 (bottom two rows), which do not co-localize with ICP4 or ICP8. Cellular DNA was stained with DAPI (blue). **E,F.** SLTM (E) and YTHDC1 (F) nuclear cv (mean±s.e.m) in noninfected (red) and infected (blue) cells following wild-type HSV1 adsorption. **G,H.** nuclear cv (mean±s.e.m) of SLTM (G) and YTHDC1 (H) in non-infected (red) and infected (blue) cells following infection by a mutant HSV1 that does not express ICP0.

The re-distribution of YTHDC1 and SLTM could be a result of their recruitment to viral replication centers. To test this we fixed infected cells six hours after infection and stained them for either ICP4 (an immediate early protein that is required for HSV1 gene expression) or ICP8 (an early protein required for HSV1 genomic replication) and found that the nuclear foci of SLTM and YTHDC1 do not co-localize with them (Fig. 6F). In fact, in agreement with the fast kinetics of the appearance of these foci, we occasionally observed cells that contained such foci but were negative for ICP8, suggesting that the re-distribution happens before viral DNA replication and is mediated by one of the immediate-early proteins of the virus.

As ICP0 is known to interact with many of the host proteins, we tested whether it is also involved in the re-distribution of SLTM and YTHDC1. Indeed, cells successfully infected with the mutant virus that does not express ICP0 did not show re-distribution of both proteins (Fig. 6G,H and Supplementary Movies 8,9).

Our results suggest that the re-distribution of SLTM and YTHDC1 into nuclear foci is an active, virus-induced process, which is facilitated by the immediate-early protein ICP0.

## DISCUSSION

Following tens of thousands of individual cells throughout the infection process we find that the outcome of HSV1 infection is largely determined by the cellular state at the time of infection. The cell's texture, morphology and cell-cycle stage at the time of adsorption enabled a supervised machine learning algorithm to predict which of the cells will become successfully infected during the next 9 hours. This variability in susceptibility among single cells is present in the population prior to meeting the virus. We find that the cellular state that makes cells susceptible to infection is composed of a fast, cell-cycle dependent component and a more stable, cell-cycle independent component. We conclude that HSV1 infection outcome is not an intrinsically stochastic event. Rather, it seems that individual cells have specific prior tendencies to become successfully-infected, showing that cellular heterogeneity of the host can have a profound impact on its survival.

We found that cell velocity is correlated with the probability of successful infection. Cell movement requires the generation of membrane extensions called filopodia (Gupton and Gertler, 2007), which several viruses have been reported to “surf” as a mechanism to enter their host cells (reviewed in (Taylor et al., 2011)). Thus, increased cell movement might be linked to a higher chance of a virus to catch a filopodia and enter the cell. An alternative explanation might be that motile cells sample more of their environment during the 30 minutes of virus adsorption, thus increasing the chances of a virus-cell meeting taking place.

Contrary to our findings, previous work done on HSV1 infection in cell populations concluded that the cell-cycle position does not affect infection outcome (Cai and Schaffer, 1991; Cohen et al., 1971). One possible explanation for this discrepancy is our enhanced sensitivity in identifying infected cells. Previous work relied on the traditional plaque-assay to measure infectivity. In this assay, 10-fold serial dilutions from a viral stock are applied to cell monolayers, which are then overlaid with agarose and infectivity is monitored several days later by manual counting of the resulting plaques in the monolayer. The noise in such measurements is often on the scale of the effect which we have measured here, of 2-3 fold change in infectivity.

In addition to its role in determining infection outcome, the cell-cycle stage also affects the speed in which infection proceeds. Cells at the earlier part of the cell cycle (G1 and early S) had, on average, a shorter lag time between viral adsorption and CFP expression than cells infected in the later part of the cell-cycle (late S and G2/M). The fact that cells at the G1/S phase are infected faster than cells in later stages might explain why HSV1 infection interferes with the natural progress of the cell-cycle, arresting cells at either the G1/S or G2/M checkpoints (Hobbs and DeLuca, 1999; Lomonte and Everett, 1999; Ehmann et al., 2000; Song et al., 2000; Paladino et al., 2014).

This finding is interesting in the context of development of new anti-viral drugs. In most cases libraries of potential compounds are assessed based on their ability to decrease the percentage of infected cells at a given time point (what we refer to as infection outcome). Such screens are likely to miss potential drugs that target infection kinetics. Infection kinetics may have a dramatic effect *in vivo*, where viral replication and dissemination is in a race against the host's mounting anti-viral response. In this context, identifying the cellular determinates of infection outcome as well as its kinetics are likely to offer new targets for drug design.

We find that the distribution of primary infection kinetics is well-described by a Gamma distribution. The Gamma distribution is defined by two parameters - rate (β) and shape (α). We find that the rate parameter varied by up to 22% between cell-cycle stages and when changing the effective moi, reflecting a scaling of the infection kinetics by these features. The shape parameter, in contrast, remained almost constant at a value of α=6 (changing by less than 5%). A Gamma distribution may arise as a result of a sequence of rate-limiting exponential processes. When this is the case, the shape parameter is the number of processes and the rate parameter is their rate (Floyd et al., 2008; Gardner et al., 2011). In the case of HSV1 infection, this may relate to a series of processes that are required for successful infection. A possible list of six of the slowest processes (as faster ones are not rate limiting) is: (1) adsorption to the cell membrane, (2) internalization, (3) binding to microtubuli, (4) binding to the nuclear pore complex, (5) mRNA synthesis and (6) protein synthesis. The measured timescale of these processes is quite similar, in the range of 15-60 minutes (Koyama and Uchida, 1987; Sodeik et al., 1997; Willis et al., 1998; Lee et al., 2016), possibly leading to the observed Gamma distribution of infection lag times, with a mean delay time of several hours.

Looking at the individual proteins in our screen, we find two proteins whose concentration at time zero is indicative of infection outcome. The concentration of one of these proteins, Geminin, is well-known to be cell-cycle regulated. In fact, the Geminin protein is part of the widely-used FUCCI system to monitor cell-cycle progression in time-lapse microscopy (Sakaue-Sawano et al., 2008). The other protein, RFX7, was not previously known to be cell-cycle regulated, but our results clearly show that its dynamics through the cell-cycle is identical to that of Geminin. Thus, the lower concentration of Geminin and RFX7 in cells that will become infected reflects the higher susceptibility of cells in the early part of the cell cycle to become successfully infected.

We did not find any other protein whose concentration or cellular localization at time zero are indicative of infection outcome. However, this should not be interpreted to suggest that such proteins do not exist. Specifically, our library does not include known anti-viral effectors, such as proteins in the NFkB and interferon pathways, because these have low expression under the basal conditions in which the library was established. Future studies looking into the effect of heterogeneity in such proteins on viral infection outcome are likely to better shape our understanding of the interplay between host and virus.

Four proteins responded specifically in successfully infected cells. The degradation of one of these proteins, SUMO2, has been previously observed and reported by others (Sahin et al., 2014; Sloan et al., 2015). The three other proteins; RPAP3, YTHDC1 and SLTM have not been previously described in the context of HSV1 infection. Interestingly, all three of these proteins seem to be related to the life-cycle of mRNA.

RPAP3 (RNA polymerase II associated protein 3) is one of the four components of the R2TP complex (von Morgen et al., 2015), a complex responsible for the assembly of several cellular machineries, including the RNA polymerase II complex. The rapid degradation of RPAP3 is in line with the previous reported alterations in RNA polymerase II upon HSV1 infection (Rice et al., 1994; Spencer et al., 1997; Jenkins and Spencer, 2001; Abrisch et al., 2015) and the known function of the virus host shutoff protein (vhs) that arrests the host's translation by degrading mRNA. The rapid block of cellular transcription and translation is likely to be important for hampering the innate-immune response in the infected cells, as the vhs protein has been shown to attenuate the host's anti-viral response (Pasieka et al., 2008; Zenner et al., 2013).

SLTM (SAFB-like Transcription modulator) is a scarcely studied homolog of SAF-B (Scaffold Attachment Factor B). Overexpression of SLTM in HeLa cells resulted in transcriptional repression and cell death (Chan et al., 2007). SAF-B is involved in the spatial arrangement of chromatin loops, poising them for transcription (Nayler et al., 1998). It can also directly bind to RNA and was shown to participate in the alternative splicing of different genes (Rivers et al., 2015). YTHDC1 is also a regulator of alternative splicing, which specifically recognizes and binds N^6^-methyladenosine (m^6^A)-containing RNAs (Xiao et al., 2016). YTHDC1 was shown to physically interact with SAF-B (Nayler et al., 1998) and all three proteins (YTHDC1, SLTM and SAF-B) were found to bind to the Xist long-non coding RNA that is required for X-chromosome inactivation (Chu et al., 2015). We find that upon HSV1 infection, SLTM and YTHDC1 change localization, forming several nuclear foci. A similar observation was made for SAF-B upon heat-shock treatment (Weighardt et al., 1999). However, in the context of HSV1 infection, the re-distribution of YTHDC1 and SLTM to nuclear foci seems to be actively controlled by the virus, as it requires the expression of the immediate-early protein ICP0.

One possible role for the sequestration of these proteins by HSV1 could be the repression of gene splicing. Unlike the majority of viruses that replicate in the nucleus, HSV1 genome contains very few introns (only 5 genes out of the ~80 encoded by the virus) and thus requires very little splicing activity. Indeed, several works have indicated that HSV1 actively represses splicing in the host (Hardy and Sandri-Goldin, 1994; Lindberg and Kreivi, 2002; Sciabica et al., 2003). This however has been called into question by a recent study that found that HSV1 causes widespread disruption of host transcription termination and that ensuing read-in into neighboring genes is responsible for the apparent inhibition of splicing (Rutkowski et al., 2015). In this regard it is interesting to note that m^6^A modification of mRNA (which is recognized by YTHDC1) is most abundant near transcriptional termination sites (Dominissini et al., 2012; Meyer et al., 2012). Whether the re-distribution of YTHDC1 and SLTM is linked to changes in the host alternative splicing or aberrant transcription termination is an intriguing question that requires further exploration.

In conclusion, this first application of the dynamic proteomics approach to study virus-host interaction at the single cell level has provided several novel insights. It allowed us to measure and model the kinetics of viral infection, to uncover the role of the host cell state in determining infection outcome and to identify new proteins that participate in the infection process. Future studies should focus on identifying the underlying molecular mechanisms that determine infection outcome and kinetics, thus opening the door to designing new and better anti-viral interventions.

## METHODS

### Library of annotated clones

The generation of the library of annotated clones (LARC) was described elsewhere (Sigal et al., 2007). The library consists of over 1,000 clones of the non-small cell lung carcinoma cell-line, H1299. All clones share the constitutively expressed mCherry-tagged proteins. The mCherry signal is bright in the nucleus and dim in the cytoplasm and is used for the automated segmentation and tracking of the cells during the analysis of time-lapse movies. Each of the clones in the library also expresses a unique YFP-tagged, full length protein. Tagging was done by the CD-tagging scheme, such that one copy of the gene is tagged in its native chromosomal location, and is thus under the control of its endogenous promoter. For all the clones in the library, the YFP-tagged protein shows a correct sub-cellular localization (clones that did not show correct localization were discarded). Cells were grown in RPMI supplemented with penicillin, streptomycin and 10% fetal bovine serum at 37C and 8% CO_2_. Cells were regularly tested for mycoplasma.

### CFP expressing HSV1

HSV-1 Strain 17 was genetically modified to express mTurq2 (a brighter derivative of CFP) from the CMV immediate early promoter by homologous recombination. The reporter gene was inserted between UL37 and UL38 genes in the viral genome, a site known to tolerate insertions with minimal effect on the viral replication. Co-transfection of purified viral DNA (Szpara et al., 2011) and plasmid pMPT06 (a kind gift from Matthew Taylor, Montana State University) was followed by several rounds of plaque purification, as described in Criddle et al. (Criddle et al., 2016).

The ICP0 mutant express mTurq2 is based was constructed using the HSV-1 dl1403 (Stow and Stow, 1986), an HSV-1 strain 17 with a 2kb deletion in both copies of the ICP0 gene (a kind gift from Roger Everett, University of Glasgow Centre for Virus Research). A viral construct originating from the mTurq2 expressing virus described above was crossed with the HSV-1 dl1403 and viral progeny where purified to obtain an ICP0 mutant express mTurq2. The progeny virus was plaque purified and tested by phenotype and by PCR to contain both the mTurq2 gene and the ICP0 deletion.

### HSV1 titration and infection

CFP expressing HSV1 was titrated in Vero cells using the Plaque assay. We assessed the infectivity of HSV1 on H1299 cells by determining the percentage of CFP positive cells at 9 hours post adsorption when infecting with serial dilutions starting at an moi of 10. We compared the observed percentage of CFP positive cells to that expected from the moi, and found that H1299 cells are approximately 10-fold less susceptible to infection than Vero cells. We used this to determine the moi for all experiments. Thus, an moi of 0.5 when infecting H1299 cells is equivalent to an moi of 5 when infecting Vero cells.

### Cells plating, infection and imaging

Cells were plated and imaged in 12-well, glass-bottom plates (MatTek, MA, US). One hour before plating the cells, plates were coated with 200µl of 10 µg/ml fibronectin from bovine serum (Sigma, Israel). Plates were washed once with 1 ml PBS and 10^5^ cells were plated in each well. Cells were allowed to grow for 24 hours. The following day, medium was replaced to an imaging medium (transparent RPMI without phenol red and riboflavin from Biological Industries, Israel, supplemented with penicillin, streptomycin and 5% fetal bovine serum) approximately one hour before infection. Medium was then aspirated and 300 µl of imaging medium containing HSV1 at an moi of 0.5 was added. Virus was allowed to adsorb to the cells for 30 minutes at 37C. During this time, the imaging set-up was performed - calibrating the microscopes, choosing four fields of view for each well and setting the acquisition times for the fluorescent channels. After 30 minutes the virus-containing medium was aspirated and 2 ml of imaging medium added to each well. Plates were placed in a temperature, CO_2_ and humidity control chambers in the microscopes, focus adjusted and imaging started. Imaging was done using two inverted epi-fluorescent Leica microscopes (DMIRE2 and DMI6000b), controled by macro scripts developed in house.

### Image and data analysis

Flat field correction and background subtraction were done for all images prior to starting the analysis. Cell segmentation and tracking was done using the PhenoTrack package for Matlab, previously developed in our lab (Sigal et al., 2006a), with some additions and modifications. All codes used in this work are available upon request. The package was extended to extract morphological and textural features. The Harlick texture features (Haralick RM, Shanmuga K, Dinstein I, 1973) were calculated using the GLCM_features1 function (http://www.mathworks.com/matlabcentral/fileexchange/22187-glcm-texture-features). Features were extracted for each segmented cell and nucleus, once from the phase channel and once from the mCherry channel. We retained the original features and also calculated the z-scored features (normalizing for all the cells in a specific clone). We also calculated the change in these features between two consecutive frames.

The CFP concentration was calculated as the median value of CFP in the cell nucleus. A threshold was calculated for each clone, based on the median level of CFP in all cells of that clone in the first five frames of the movie (less than two hours post HSV1 adsorption).

To ensure correct tracking of the cells we employed several filters, eliminating trajectories of cells that did not meet certain criteria. Such criteria included, for example, more than one mitosis event in 12 hours and a rapid, non-physiological, change in the mCherry levels. Overall we eliminated ~2/3 of the data, remaining with ~52,000 reliably tracked cells out of ~190,000 cells imaged in total.

### Supervised machine learning for predicting infection outcome

We divided our dataset of ~52,000 cells into two group - infected (CFP positive at 9 hours post HSV1 adsorption) and non-infected (CFP negative for the entire 12 hours). Next we divided the data into train and test sets. To avoid any biases due to differences between the clones, we made sure that each clone is similarly represented in the infected and non-infected groups. We additionally made sure that no particular clone will be over represented in the dataset. 75% of the data was used for training the classifier and 25% (from clones not used in the training step) for testing its performance. The training set included 23,780 cells and the test set 8,108.

We used Matlab version R2015b for all supervised machine learning procedures. We used Matlab's *fitensemble* function to construct decision trees for classification using the RobustBoost algorithm. We performed feature selection by identifying the 20 features with highest predictive power using the *predictorImportance* function. The final classifier included 2,000 decision trees based on the top 20 features.

### Extracting cell-cycle data from still images

We employed a supervised machine learning approach, similar to that used by others (Kafri et al., 2013; Gut et al., 2015; Blasi et al., 2016), which infers the cell-cycle position of a cell from a still image using a random forest regression predictor. The performance of this predictor is shown in Supplementary Fig. 3. We trained and tested the predictor using independent datasets of non-infected cells that divided during the movies, so that we could determine the time after mitosis for each cell in each frame. We aligned the cells trajectories to an imaginary cell-cycle length of 24 hours. This gave the best results, but using other cell-cycle lengths did not significantly alter our findings. We selected the top 30 features to use in the predictor.

### Infection kinetics modeling

We fitted the distribution of infection lag times with a three-parameter mixture model for the primary and secondary infections (Supplementary Fig. 2). The model assumes that the lag time between adsorption and infection can be captured by a two-parameter distribution, and that this distribution also captures the lag between primary and secondary infections. Specifically, the number of secondary infection at a given time-point depends on the number of infections in all previous time-points with appropriate delays that are given by the two-parameter distribution. The relative number of secondary infections is also fitted as a third parameter. Overall we fitted three parameters - two for the distribution (*α,β*) and one for the relative number of secondary infections (*δ*).

We fitted the following two parameter distributions: Normal, Log-Normal, Weibull and Gamma. For each distribution we scanned each parameter with a resolution of 0.05. Each parameter was scanned in the range of ±1 of the best fit value and *δ* was scanned in the range of 0-2. For each set of parameters we generated a distribution from the mixture model and estimated the log-likelihood of the data given this distribution.

To statistically assess which distribution fits the data best we performed a bootstrapping procedure. We generated a distribution of log-likelihoods for each fit by resampling our data 1,000 times with replacements. We then performed a one-sided t-test to compute the significance of the difference between these distributions. The Gamma distribution fitted our data significantly better than the other three (maximal p-value<10^−15^).

Confidence intervals for each parameter were computed by fixing other parameters and fitting a third order polynomial to the distribution of log-likelihoods around the fitted parameter. We then computed the 95% confidence interval using the second derivative of this polynomial at this parameter. For all the parameters assessed, confidence intervals were at least one order of magnitude lower than the estimated parameter.

### Cell-cycle synchronization

5X10^4^ cells were plated in 12-well glass bottom plates as described above. At 5pm the medium was replaced with a full medium containing 2 mM thymidine (Sigma-Aldrich, Israel). At 8am the next morning cells were washed twice with PBS and normal growth medium was added. At 5pm of the same day the medium was again replaced with a thymidine containing medium. At 8am the next morning half of the wells were released from blocking (washed twice and given normal growth medium) and half were maintained in the blocking medium. At 4pm, eight hours later, the blocked cells were released. Cells were washed and infected with HSV1 at an moi of 0.25 or 0.5 and imaged as described above.

One well from each condition (15 minutes or 8 hours post-release) was harvested and fixed with 70% ethanol at the time of HSV1 infection. Samples were rehydrated, treated with RNAse A and stained with propidium iodide to analyze the cell-cycle stage by FACS.

## ACKNOWLEDGMENTS

This work has been supported by an Israeli Science Foundation (ISF) grant from the Israeli Academy of Sciences (grant number #1349/15). ND wishes to thank the Clore Foundation for their generous support through the Sir Charles Clore Postdoctoral fellowships program. We thank Prof Matthew D. Weitzman for helpful discussions and critical reading of the manuscript.

---

**Supplementary Figure 1.**
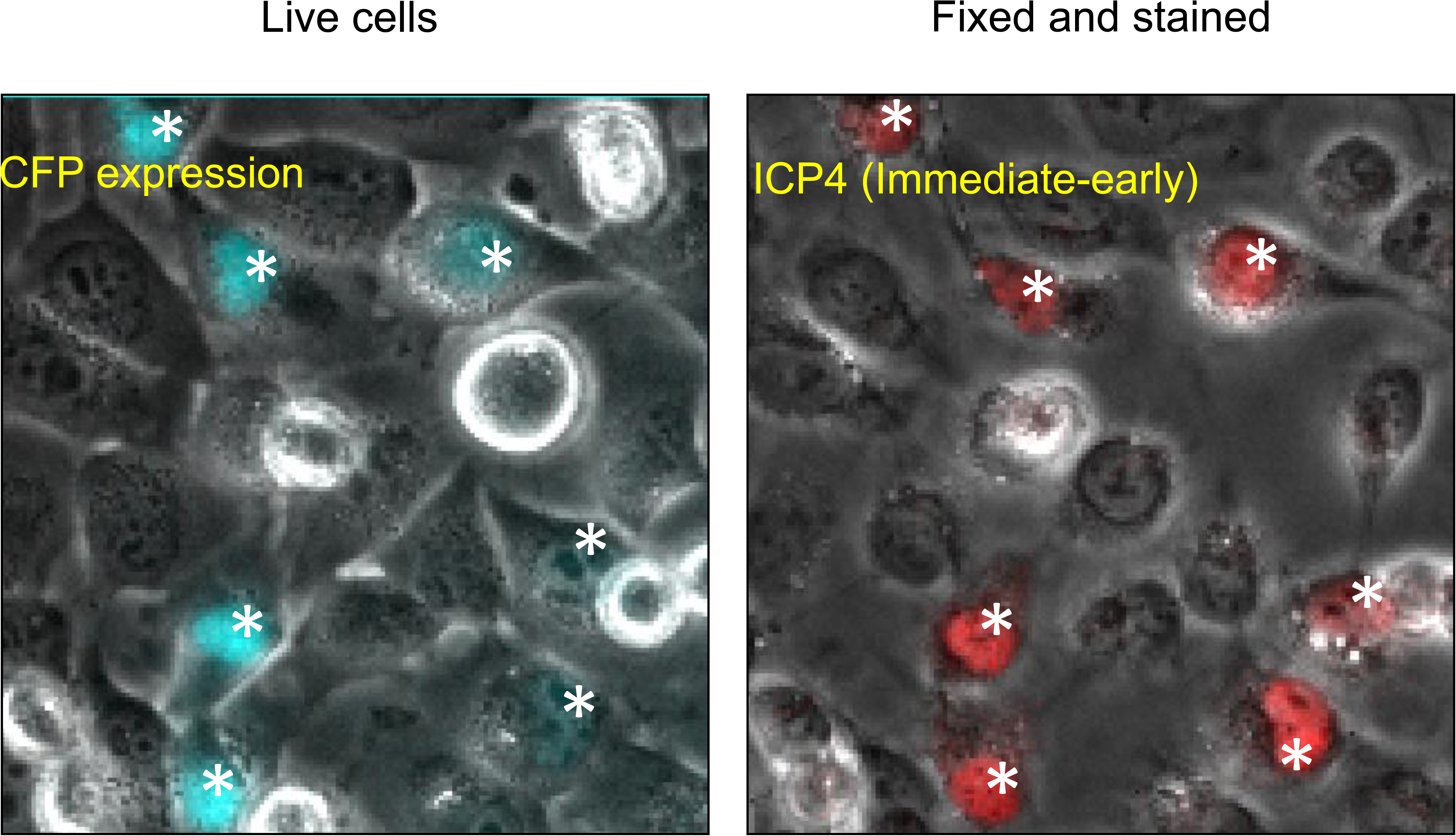
CFP expression correlates with HSV1 immediate-early protein expression. Cells infected with HSV1 expressing CFP were imaged at 7 hours post adsorption, fixed and stained with antibodies against different viral proteins. All the CFP positive cells are also positive for ICP4, an immediate-early protein that is synthesized before viral DNA replication. Asterisks mark CFP positive cells and the corresponding cells after fixing and staining.

**Supplementary Figure 2.**
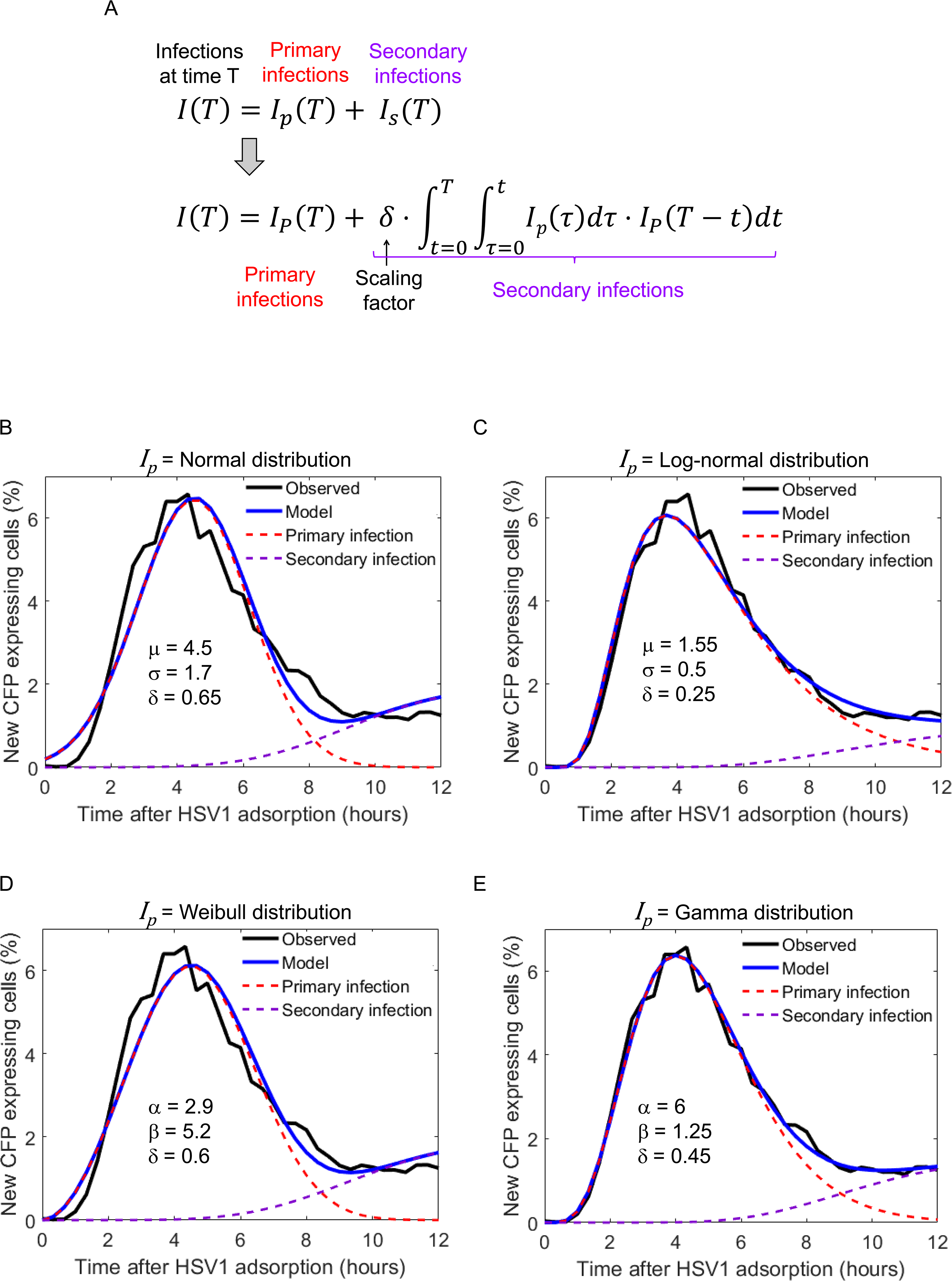
kinetics of primary infection are best described by a Gamma distribution. **A.** Modeling infection kinetics as the sum of primary and secondary infections (top row). We assume that the secondary infections have the same delay kinetics as primary infections, and thus can be rewritten as a function of the primary infection distribution, *I*_*p*_ (bottom row). **B,C,D,E**. We estimated the fit for *I*_*p*_ using four distribution types – Normal (B), Log-normal (C), Weibull (D) and Gamma (E). These four distribution types are defined by two parameters. The best fitting parameters are presented in the figures. In addition we also fitted the scaling factor δ. Of these four distribution, Gamma gave the best fit the observed data (black lines). The blue lines show the model with best parameters, which can be further divided into primary (dashed red lines) and secondary (dashed purple lines) infections.

**Supplementary Figure 3.**
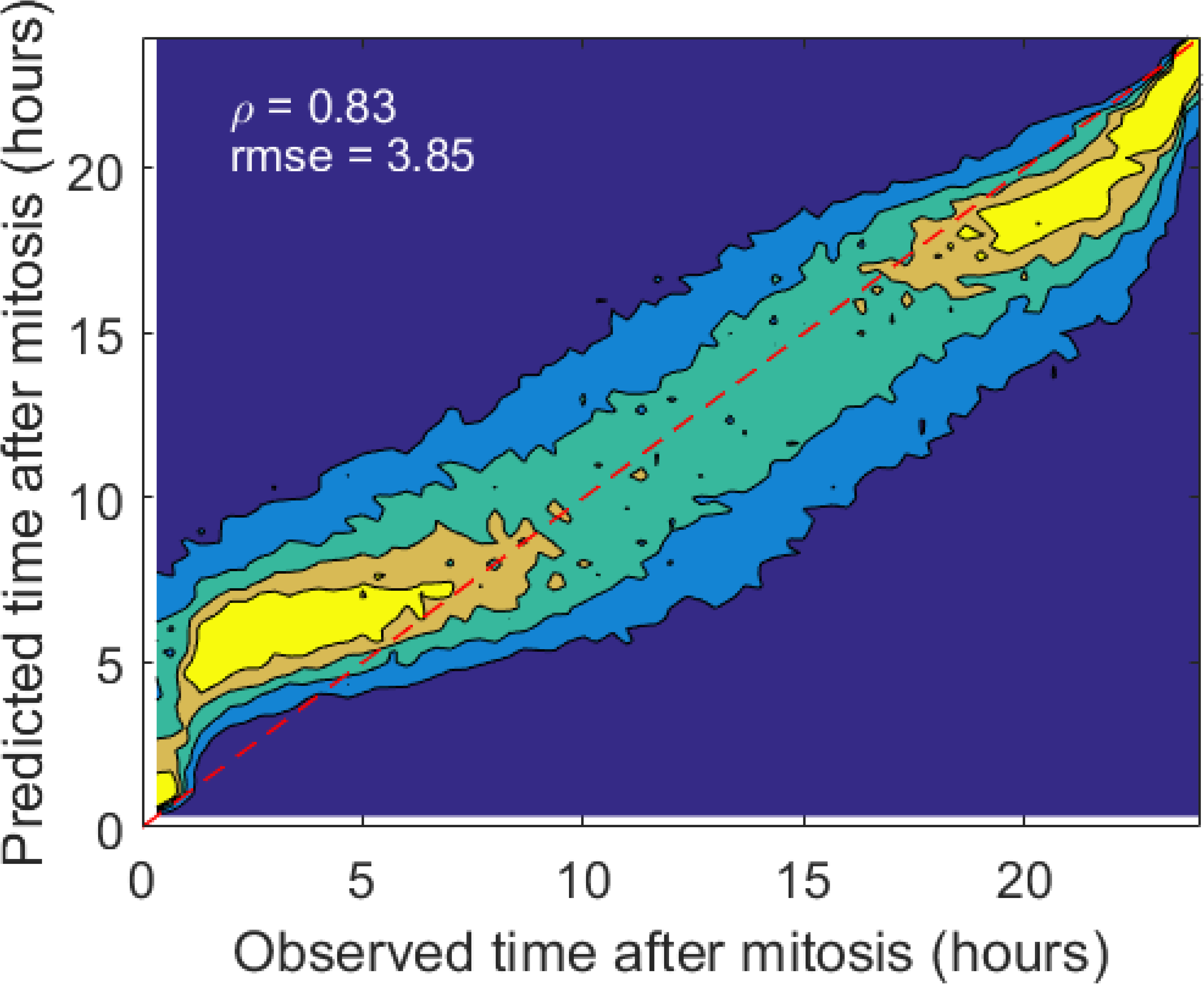
Cell cycle prediction from still images and cell-cycle effect on HSV1 infection outcome. To asses the cell-cycle stage of the cells at the time of infection we trained a random forest predictor, using a dataset of noninfected cells that divided during the time-lapse movies. We used the top 30 predictive features and trained an ensemble of 500 decision trees. The figure shows the performance of the predictor on an independent test set. The predictor calculates the time from last mitosis with an rmse of 3.85. the Pearson correlation coefficient was 0.83.

**Supplementary Figure 4.**
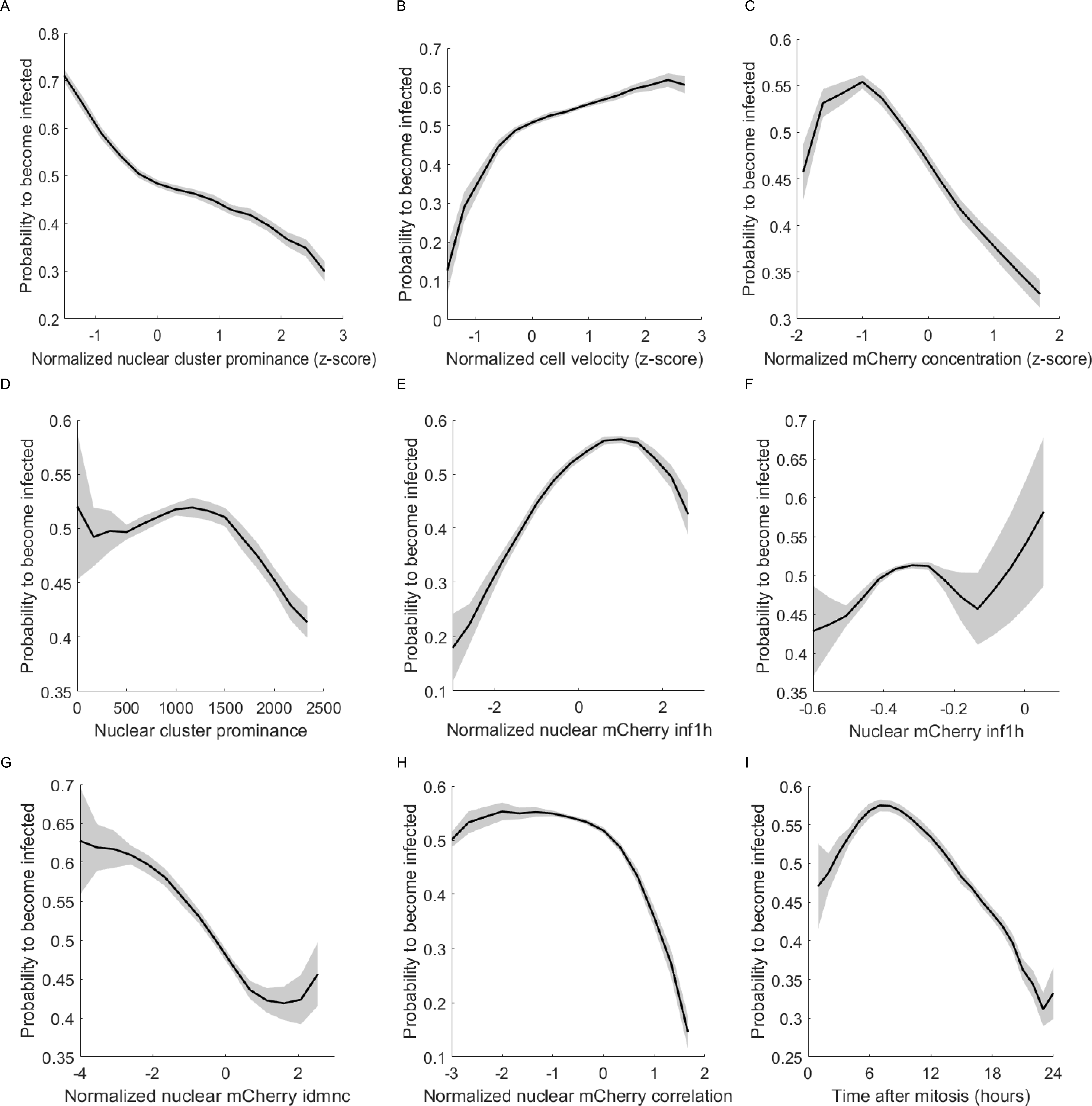
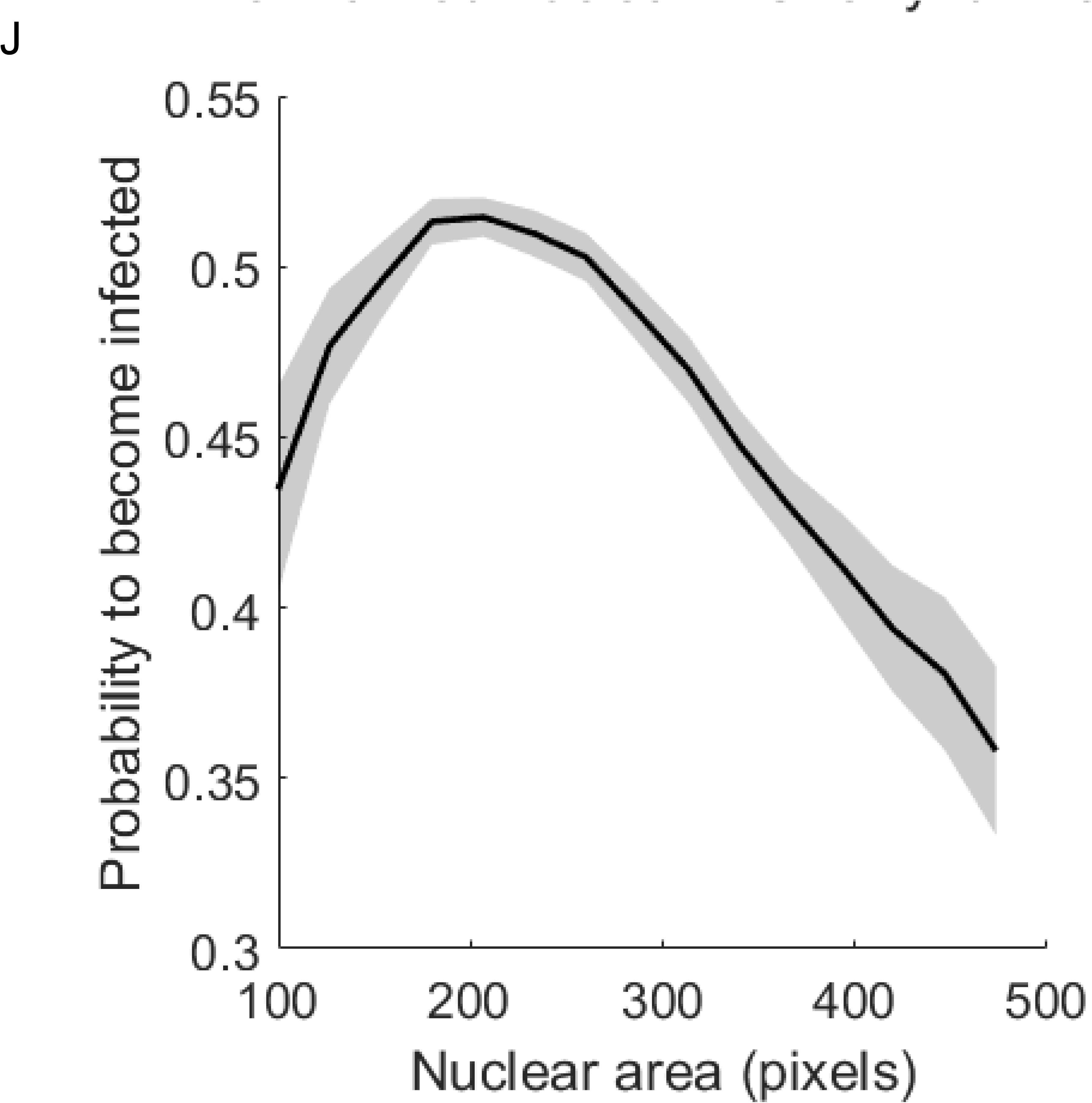
Infection probability as a function of different features. We analyzed the relation between infection probability and the top 10 features used by the supervised machine learning algorithm to predict infection outcome. Infection probability was calculated by binning the cells and calculating the percentage of infected cells in each bin. The features are: (A) normalized cluster prominence of the nucleus from the phase channel, (B) normalized cell velocity, (C) normalized mCherry concentration, (D), raw cluster prominence of the nucleus from the phase channel, (E) normalized information measure of correlation 1 of the nucleus from the mCherry channel, (F) raw information measure of correlation 1 of the nucleus from the mCherry channel, (G) normalized Inverse difference moment normalized of the nucleus from the mCherry channel, (H) normalized correlation of the nucleus from the mCherry channel, (I) cell-cycle stage and (J) nuclear area

**Supplementary Figure 5.**
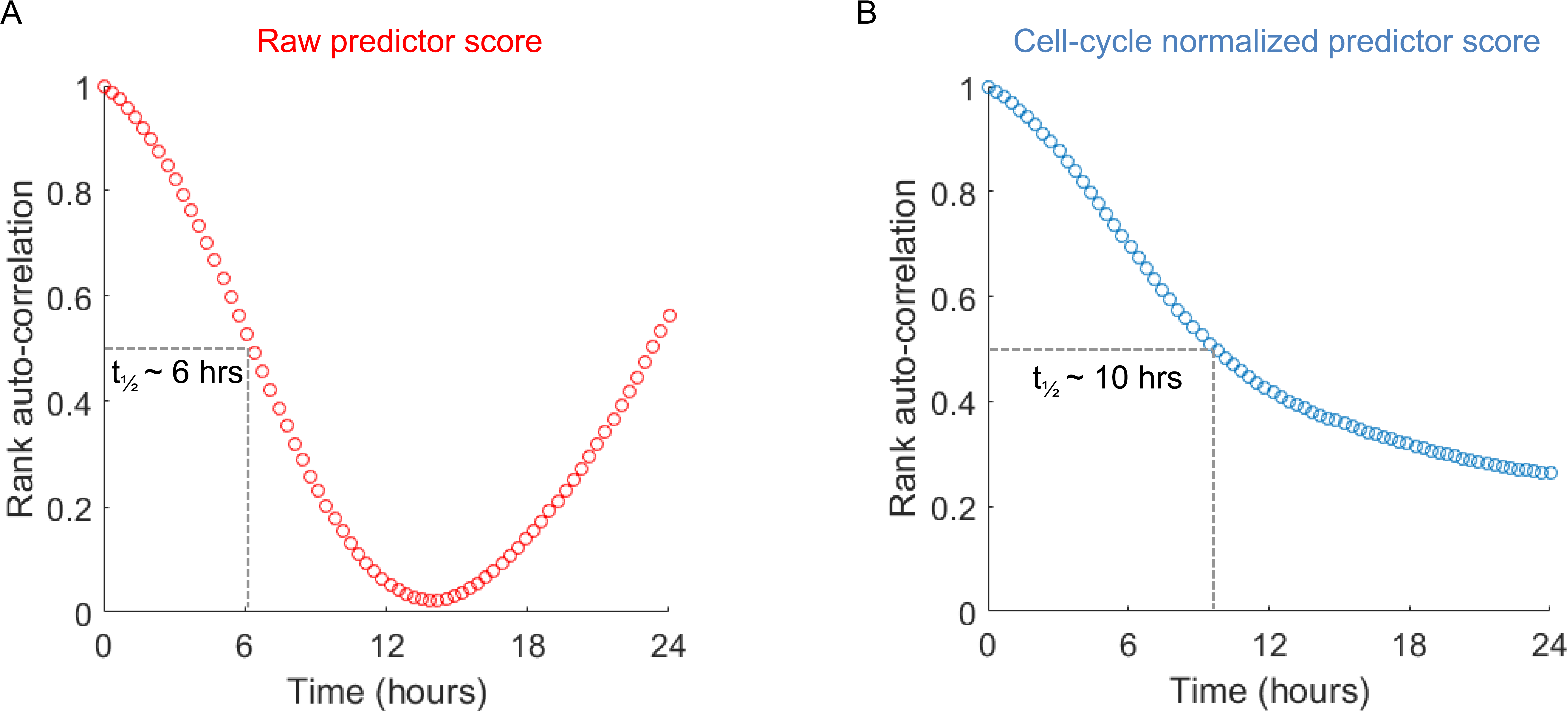
Rank auto-correlation of predictor score in individual cells over time. Predictor score were calculated for >200 cells that were tracked for 24 hours prior to infection. The cells were ranked according to their score in the first frame, and the rank auto-correlation was calculated. Shown are the auto-correlation for the raw predictor score (A) and after normalizing the score according to the cell-cycle position of the cell (B). The raw scores decays faster, with a t_1/2_ time of about 6 hours. It reaches zero around half a cell-cycle (12 hours) and then begins to raise again. After normalizing for the cell-cycle effect, the auto-correlation decay is slower, with a t_1/2_ around 10 hours.

